# Parallel cortical-brainstem pathways to attentional analgesia

**DOI:** 10.1101/2020.02.20.955161

**Authors:** Valeria Oliva, Rob Gregory, Wendy-Elizabeth Davies, Lee Harrison, Rosalyn Moran, Anthony E. Pickering, Jonathan C.W. Brooks

## Abstract

Pain perception is diminished when attention is diverted. Our previous human fMRI study, using a 2×2 factorial design with thermal stimuli and concurrent visual attention task, linked the brainstem triad of locus coeruleus (LC), rostroventromedial medulla (RVM) and periaqueductal grey (PAG) to attentional analgesia. This study was repeated with a larger cohort, replicating our earlier findings. Pain intensity was encoded by the RVM, whilst activity in the contralateral LC correlated with the magnitude of attentional analgesia. Psycho-Physiological Interaction analysis informed subsequent Dynamic Causal Modelling and identified two parallel paths between forebrain and the brainstem regions involved in analgesia. These cortico-brainstem connections were modulated by attentional demand: a bidirectional anterior cingulate cortex (ACC) – right-LC loop, and a top-down influence of task on ACC-PAG-RVM. Under conditions of competing attentional demands the ACC recruits discrete brainstem circuits to modulate nociceptive input.

## Introduction

Attentional analgesia is a well-characterised phenomenon whereby increased cognitive load can decrease pain perception (Peyron et al. 2000; Bantick et al. 2002; Brooks et al. 2002; Valet et al. 2004; Brooks et al. 2017; Sprenger et al. 2012). For example, this can be achieved by diverting attention from a painful stimulus to a visual task or simply by active mind-wandering (Bushnell et al., 2013; Kucyi et al., 2013). Central to attentional analgesia is the concept of divided attention, whereby less cognitive resource is available to be allocated to nociception and pain. Since noxious stimuli are inherently salient and therefore attention grabbing (Eccleston et al., 1999), then any concurrent cognitive task must compete for ‘attentional’ resource. Attention is thus cast both as a key component of pain behaviour (i.e. attending to pain (Crombez et al., 2004; Legrain et al., 2009; Roelofs et al., 2002)) as well as a putative mechanism for pain relief. The processes regulating attentional focus is of importance in the development, maintenance and potentially resolution of chronic pain states.

The mechanisms that allow attention to regulate pain are currently not well understood and there has been ongoing debate about whether attentional analgesia requires engagement of descending control to attenuate nociception (Brooks et al., 2017; Bushnell et al., 2013; Lorenz, Minoshima, & Casey, 2003; Tracey et al., 2002; Valet et al., 2004). Several important regions have been linked to the descending analgesic effects, including the anterior cingulate cortex (ACC), dorsolateral prefrontal cortex (dlPFC) and components of the descending pain control system including periaqueductal grey (PAG), rostroventromedial medulla (RVM) and locus coeruleus (LC). An interaction between cortical and mid-brain structures during distraction from pain has been identified (Lorenz et al., 2003; Valet et al., 2004) but these previous studies were unable to examine interactions in pontomedullary regions that are known to be important for the descending control of nociception.

These brainstem regions are all candidates for mediating attentional analgesia given their known anti-nociceptive roles (Millan, 2002). For example, multiple animal studies have demonstrated that interactions between the PAG and RVM produces endogenous analgesia, mediated by spinally projecting neurons in the RVM (Basbaum et al., 1979; Fields et al., 1978; Heinricher et al., 2009). Together with the ACC, these regions form one of the main pain modulatory pathways involved in the bidirectional modulation (i.e. facilitation and inhibition) of nociception in the spinal cord dorsal horn (De Felice et al., 2016; Ossipov et al., 2010; Quintero, 2013).

Similarly, the LC is another potential candidate region that could mediate the interaction between attention and pain because of its projections to the spinal cord which release noradrenaline to produce analgesia (Hirschberg et al., 2017; Llorca-Torralba et al., 2016). Additionally, it has a known role in salience signalling and attention mediated by ascending projections (Aston-Jones et al., 1999; Sales et al., 2019; Sara et al., 2012). Despite it being challenging to resolve with fMRI (Astafiev et al., 2010; Liu et al., 2017), the LC was recently identified as the only region whose activity reflected the interaction between task and temperature in an attentional analgesia paradigm (Brooks et al., 2017). The LC could therefore contribute to attentional analgesia as part of the PAG-RVM system, or as a parallel descending modulatory pathway perhaps receiving inputs directly from ACC (Aston-Jones et al., 1991; Bajic et al., 1999).

Within this framework the ACC is ideally placed to mediate between competing cognitive demands (e.g. between a sustained visual attention task and pain) as it is active during conflict resolution (Braver et al., 2001; Kerns, 2006; Kim et al., 2011), its activity is modulated by attention (Davis et al., 2000) as well as being consistently activated by painful stimuli (Brooks et al., 2017; Garcia-Larrea et al., 2013; Peyron et al., 2000; Wager et al., 2013). The ACC is known to code for pain intensity (Büchel et al., 2002; Coghill et al., 2003) and unpleasantness (Rainville et al., 1997), furthermore, sub-divisions (e.g. dorsal anterior ACC) are involved in high level cognitive appraisal of pain, including attention (Büchel et al., 2002). Some have proposed a specific role for dorsal ACC (dACC) in pain perception (Lieberman et al., 2015), though this is disputed with other studies suggesting that activity within this structure reflects the multifaceted nature of pain (Wager et al., 2016). Connectivity between the ACC and structures involved in descending pain control e.g. the PAG, has been shown to vary with pain perception due to both attentional modulation of pain and placebo analgesic responses (Bantick et al., 2002; Eippert et al., 2009; Petrovic et al., 2000; Valet et al., 2004) suggestive of a role in attentional analgesia. Indeed, cingulotomy has been shown to “disinhibit” noxious heat and cold perception leading to hyperalgesia (Davis et al., 1994).

We hypothesised a top-down pathway mediating attentional analgesia where the PAG receives attentional-shift signals from the ACC and/or LC and directs the RVM and/or LC to attenuate nociceptive processing in the spinal cord. Given the multiplicity of possible pathways and interactions by which activity in the brainstem can generate analgesia, we anticipated that effective connectivity analyses could resolve the roles of these regions (identified in our previous investigation (Brooks et al., 2017)) during attentional analgesia. To increase the statistical power to undertake this connectivity analysis, two further fMRI datasets were acquired using the same paradigm as per Brooks et al. (2017). Analysis of these additional datasets reproduced our previous regional activation results and so the three datasets were pooled for the effective connectivity analyses and modelling. We tested for psycho-physiological interactions (PPI, (Friston et al., 1997; McLaren et al., 2012; O’Reilly et al., 2012) to explore whether the connectivity between the PAG, RVM, LC and ACC altered during the experimental paradigm. Finally, we used dynamic causal modelling (DCM, Friston et al. 2003) to test the directionality and strength of the connections.

## Methods

### Participants

Subjects were recruited using poster and email adverts at the University of Bristol for three different pain imaging studies at the Clinical Research and Imaging Centre (CRiCBristol) that used the same experimental paradigm: an initial study on attentional analgesia (Brooks et al., 2017), a study on sleep disruption and a study on fibromyalgia. The first two studies were approved by the University Bristol, Faculty of Science, Human Research Ethics Committee (reference 280612567 and 291112606 respectively) and the fibromyalgia study was approved by NHS South Central Oxford B Research Ethics Committee (reference 13/SC/0617).

All subjects gave written informed consent after application of standard inclusion/exclusion criteria for participation in MRI studies. The presence of significant medical/psychiatric disorders (including depression) or pregnancy precluded participation. Subjects with a chronic pain condition, or those who were regularly taking analgesics or psychoactive medications were also excluded. All subjects were right-handed, verified with the Edinburgh handedness inventory (Oldfield, 1971).

The *discovery cohort* were 20 right-handed healthy subjects (median age 25 years, range 18–51 years, 10 females). Subjects attended for two sessions. During the screening visit, written consent was obtained and both task difficulty and temperature of the thermal stimulation were individually calibrated. Subsequently the subjects returned for the test session where they completed the experiment in the MRI scanner (For full details on the *discovery cohort* see Brooks et al., 2017).

The *validation cohort* composed of control subjects from two separate studies:

Twenty healthy volunteers (median age 23, range 20-33, 10 females) were recruited for a study investigating the effects of sleep disturbance on attentional analgesia. Subjects completed the same experiment protocol on two occasions; after a habitual and a disturbed night’s sleep (at the sleep laboratory at CRiCBristol). For the present study, only data obtained from the control condition was used, wherein subjects experienced their habitual sleep regime the night prior to their scan.

A second group of 20 healthy participants (median age 31.5, range 20-59, 18 females) was recruited from the control group of a study analysing attentional analgesia in fibromyalgia patients.

### Experiment

Thermal stimuli were delivered to the left volar forearm (approximately C6 dermatome) using a circular contact thermode (CHEPS Pathway, MEDOC) and each lasted 30 seconds. The noxious thermal stimulus was individually titrated to obtain a 6 out of 10 pain rating (42-45°C plateau). The innocuous stimulus plateau was set at 36°C. In both cases brief heat spikes of 2, 3 and 4°C above the plateau temperature were added in a random sequence at a frequency of 1Hz. This heating profile was used to maintain painful perception, whilst avoiding skin sensitisation. The baseline thermode temperature was 32°C.

For the Rapid Serial Visual Presentation task (RSVP, Potter et al. 1969), subjects identified a visual target (the number “5”) among distractors (other letters and numbers), presented using back-projection to a screen, responding with a button box (Lumina LP-400, Cedrus). The speed of character presentation for the hard RSVP task was individually calibrated to obtain a 70% correct detection rate using d’, and ranged from 32 to 96 ms. The speed of presentation for the easy RSVP task was either 192 or 256 ms, depending on performance in the hard task (if the “hard” task interval for the subject was <80 ms or >80 ms, respectively).

### Data acquisition

In the scanner, participants received noxious or innocuous thermal stimuli (high/low) while simultaneously performing the RSVP task with two levels of difficulty (easy/hard). Thus, there were four experimental conditions (in a 2×2 factorial experimental design): *easy|low*, *easy|high*, *hard|low*, *hard|high*. Each condition was repeated 4 times. Each experimental epoch started with instructions (5s), followed by the 30s experimental condition, followed by a 10s rest period before an 8s rating period where subjects rated the perceived pain intensity from 0 to 10 on a visual analogue scale (VAS) (See Fig 1 in Brooks et al. 2017). The post-stimulus interval, between the rating period and subsequent instructions, was 17s.

The experiment for the *validation cohort* (n=38) was essentially identical to that of the *discovery cohort*. The titrated mean high temperature for the *discovery cohort* was 44.2°C and for the *validation cohort* it was 43°C (range 42–45 °C). The whole imaging session lasted 26 minutes for the *discovery cohort* and sleep-disruption cohort and was 22 minutes for the fibromyalgia cohort (as a redundant control condition, with no distraction during high temperature, was omitted).

Imaging was performed with a 3T Skyra MR system (Siemens Medical Solutions, Erlangen, Germany) and 32-channel receive only-head coil. In addition to blood oxygenation level dependent (BOLD) functional data, T1 weighted structural scans were acquired with an MPRAGE sequence to allow image registration. Functional imaging data were acquired with TE/TR=30/3000 ms, GRAPPA acceleration factor = 2, resolution = 1.5 x 1.5 x 3.5 mm. The slices were angulated perpendicularly to the base of the 4^th^ ventricle to improve the acquisition of brainstem nuclei. This slice orientation optimised the ability to discriminate between the small brainstem structures in the transverse plane and while allowing the capture of whole brain activity within 3 seconds. Fieldmap data were acquired with a gradient echo sequence (TE1/TE2/TR = 4.92 / 7.38 / 520 ms, flip angle 60°, resolution 3 x 3 x 3 mm). During scanning, a pulse oximeter and a respiratory bellows (Expression MRI Monitoring System, InVivo, Gainesville, FL) were used to monitor cardiac pulse waveform and respiratory movement for subsequent physiological noise correction (Brooks et al., 2013).

### Behavioural Data analysis

Pain VAS ratings were converted to a 0-100 scale for a repeated measures ANOVA in SPSS software (after Brooks et al. 2017). Following estimation of main effects (task, temperature) and interactions, post-hoc paired t-tests were performed. The presence of attentional analgesia was pre-defined as a significant interaction between task difficulty and high temperature on pain rating on post-hoc paired t-testing (p < 0.05). To test for differences between the *discovery* and *validation* cohorts; group membership was added as a between subject factor to the two within subject factors (task and temperature). Subsequent analysis is reported on the *pooled* cohort.

### Imaging Data analysis

#### Image Pre-processing

Functional images were corrected for motion using MCFLIRT (Jenkinson et al., 2012) and co-registered to each subject’s structural scan using brain boundary-based registration (Greve et al., 2009) and then to the 2mm template (“MNI152”) brain using a combination of fieldmap based unwarping using FUGUE (Jenkinson, 2003), linear transformation using FLIRT (Jenkinson et al., 2001) and non-linear registration using FNIRT (Andersson et al., 2007) with 5mm warp spacing. Functional data were spatially smoothed with a kernel size of 3mm (FWHM) and high pass temporally filtered with a 90s cut-off. Two subjects in the *validation* cohort and one from the *discovery cohort* were excluded from the analyses at this stage because of signal dropout in the EPI data (leaving 57 subjects).

#### First level analyses

Local autocorrelation correction was performed using FILM (Woolrich et al., 2001) as part of model estimation, which also attempted to correct for physiologically driven signals (originating from cardiac/respiratory processes) using slice-dependent regressors (Brooks et al., 2008; Harvey et al., 2008). The four conditions (*easy|high*, *hard|high*, *easy|low*, *hard|low*) and tasks of no interest (cues and rating periods) were modelled using a hemodynamic response function (gamma basis function, σ = 3s, mean lag = 6s) alongside the physiological regressors within the general linear model in FEAT (Jenkinson et al., 2012). A separate analysis tested for an intra-subject parametric relationship between pain ratings (one per block) and BOLD signal (Büchel et al., 1998). In addition to tasks of no interest and physiological signal regressors, a constant regressor for all blocks (weighting = 1) and a regressor weighting the individual pain ratings for each block were included. None of the regressors were orthogonalised with respect to any other.

#### Second level analyses

Main effects were specified as positive and negative main effect of attention (hard versus easy task, and vice versa) and positive and negative main effect of temperature (high versus low thermal stimulus, and vice versa). A task x temperature interaction contrast was also specified. The parametric data was assessed using a simple group average – to examine whether the linear relationship between pain ratings and brain activity was consistent across the group. Lastly, a paired analysis compared activity during the *easy*|*high* and *hard*|*high* conditions - to examine whether the inter-subject difference in average pain ratings (i.e. *easy|high* minus *hard|high*) was linearly related to the corresponding difference in BOLD signal (similar to Tracey et al. 2002 and Brooks et al. 2017). To test for differences between the *discovery cohort* and the *validation cohort,* we used an unpaired t test with FLAME (height threshold z > 3.09, corrected cluster extent threshold p < 0.05), in line with guidelines on corrections for familywise error (FWE) (Eklund et al., 2016). Subsequent analyses of the *pooled* cohort (i.e. all 57 subjects) used the same threshold.

#### Brainstem-specific analyses

Detecting activation in the brainstem is non-trivial due to its susceptibility to physiological noise and artefacts (Brooks et al., 2013), small size of structures of interest and relative distance from signal detectors in the head coil. Consequently, a brainstem focussed analysis was performed at the group level using a series of anatomical masks and statistical inference using permutation testing in RANDOMISE (Nichols et al., 2002). Analyses utilised pre-defined regions of interest based on (i) a whole brainstem mask derived from the probabilistic Harvard-Oxford subcortical structural atlas (Desikan et al., 2006) and thresholded at 50% and (ii) previously defined probabilistic masks of the *a priori* specified brainstem nuclei (RVM, LC, PAG) from Brooks *et al.* (2017). The number of permutations were set to 10,000 in line with guidelines (Eklund et al. 2016) and results reported using threshold free cluster enhancement (TFCE) corrected p < 0.05 (Smith et al., 2009).

#### Psycho-Physiological Interactions (PPI)

Effective connectivity analyses were performed on the *pooled cohort*. We used generalised PPI (gPPI) to detect changes in interactions between regions during specific experimental conditions (O’Reilly et al. 2012; McLaren et al. 2012; Friston et al. 1997). In this technique a physiological signal (e.g. the time-course extracted from a seed region) is convolved with a modelled psychological variable (i.e. each one of the experimental conditions) to build an interaction regressor. All interaction regressors were added to a general linear model (GLM) that also included the non-convolved experimental conditions and tasks of no interest (e.g. the rating period). Contrasts were built to test for connectivity differences that could be explained by the main effects of task and temperature and the task * temperature interaction.

Four regions identified in the main effect analyses (temperature and/or attention) in the *pooled cohort*, were selected as seed-regions for the gPPI analysis: PAG, right LC and ACC in the main effect of task and RVM in the main effect of temperature. For each subject, the physiological BOLD time course was extracted from the peak voxel of the pre-processed images (as described in the previous section) within each functional mask, and gPPI performed at the first level. Subsequently, group responses were estimated with permutation testing within the same functional masks e.g. effective connectivity between PAG seed region and the other three regions (RVM, right LC, ACC). For a post hoc investigation of significant results in the task * temperature interaction contrast, we focussed on the conditions of interest (i.e. *easy|high* and *hard|high*). Parameter estimates were extracted by first defining a sphere of radius 2mm at the voxel of greatest significance in the group gPPI result, then back-transforming this mask to subject space and extracting the signal from the voxel with highest Z-score.

In summary, the procedure for gPPI analysis was:

- Time series extraction from functional masks in pre-processed data
- Convolution of time-series with experimental condition
- Building statistical contrasts in a GLM
- First level (single subject) analysis
- Group analysis permutation testing with functional masks
- Extraction of parameter estimates from the conditions of interest.

#### Dynamic Causal Modelling (DCM)

Given the inability of gPPI to resolve the directionality of connections, we sought to extend our findings by using DCM (Friston et al., 2003). This technique allows the specification of a hypothetical network model (based on equation 1) fitted to the fMRI data to resolve connection strengths.

The change in activity of each region in a model with *j* inputs and *n* brain regions is formalized as follows:

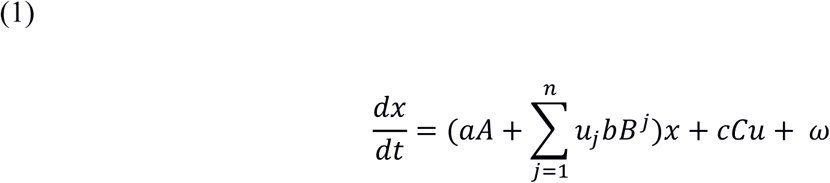

Where:

*x* - neuronal state of a region (i.e. BOLD signal convolved with haemodynamic response function)
*A* - binary vector that defines the connectivity of *x* is to each of the other regions in the model,
*a* - vector of parameters that define the strengths of such connections,
*u* - external input to the model,
*B* - binary vector that defines whether model connections are modulated by external input,
*b* - vector of parameters that defines the strength of such modulation,
*C* - binary vector that defines whether x directly receives the external input,
*c* - contains parameters that regulate the strength of the received input,
*ω* - random neuronal noise.

Note since the model is estimated in a Bayesian framework, parameters are not single values but are posterior densities.

Given the results of the PPI analysis, we specified bi-linear, one state, stochastic, input centred DCMs (Daunizeau et al., 2009; Daunizeau et al., 2012) in SPM 12 (Wellcome Trust Centre for Neuroimaging, London, UK). The models were estimated on a computer cluster (BlueCrystal) in the Advanced Computing Research Centre, University of Bristol – http://www.bristol.ac.uk/acrc/.

Random effects Bayesian Model Selection (BMS) was used to compare the models and Protected Exceedance Probability, the likelihood of a given model in respect to the others tested, was calculated. Bayesian Omnibus Risk, a measure of the risk of all models having the same frequency within the population, was also computed (Rigoux et al., 2014). Bayesian model averaging (Penny et al., 2010) was used to extract the parameter estimates of interest.

## Results

### Comparison of the discovery cohort and validation cohort

We initially analysed pain scores and main effects of task and temperature on brain activation maps in the *validation cohort* (n=38) independently to qualitatively compare against the results previously obtained in the *discovery cohort* (n=19, Brooks et al., 2017). A two-way repeated measures ANOVA on the pain ratings in the *validation cohort* revealed a main effect of temperature (F(1, 37) = 137.5, P < 0.0001) and a task x temperature interaction (F(1, 37) = 24.6, P < 0.0001, Fig. S1), and no main effect of task (F(1, 37) = 2.1, P = 0.15). A post-hoc paired t-test showed performance of the hard task produced a decrease in pain scores in the high temperature condition (mean *hard|high* =39.3, SD 18.9 vs *easy|high* =43.6, SD 18.3, P < 0.001, Bonferroni corrected), consistent with an attentional analgesic effect (Fig S1).

Analysis of the fMRI activation maps in the *validation cohort* showed a main effect of temperature (*high* versus *low* temperature) in a range of regions including the primary somatosensory cortex, dorsal posterior insula and opercular cortex, with more prominent clusters contralateral to the side of stimulation (applied to the left forearm, Fig S2). In the brainstem, permutation testing with PAG, LC and RVM masks showed activation in all of these nuclei (Fig S2). No cluster reached significance in the negative main effect of temperature (low temperature versus high temperature).

Analysis of the main effect of task (*hard* versus *easy* task) showed bilateral activation in the occipital cortex (Fig S3). The anterior cingulate, anterior insula and paracingulate cortices also showed extensive activation, consistent with the visual attention network (Wager et al., 2004). Activation in the dorsolateral PAG was observed in this whole brain analysis without the need for brainstem-specific masking (Fig S3), presumably because of the increased statistical power afforded by the higher number of subjects compared to Brooks et al. (2017). A permutation test with PAG, LC and RVM masks showed activation in all nuclei (Fig S3). In the reverse contrast (negative main effect of task, *easy* versus *hard*) main activation clusters were located in the posterior cingulate cortex, frontal medial cortex and in the lateral occipital cortex.

Activation clusters from these main effects analyses are reported in table S1 and, as indicated, were closely consistent with our previous results (Brooks et al, 2017). This qualitative comparison was also assessed quantitatively as follows:

A three-way repeated measures ANOVA on the pain scores using task and temperature as within subject factors and the group (*discovery* vs *validation cohort*) as between subject factor showed no effect of group on the effects of temperature (P = 0.481), nor task (P = 0.833), nor on the task*temperature interaction (P = 0.481).

An unpaired t-test on the functional image contrasts did not show any statistically significant differences between the *discovery* and *validation cohorts* for the main effect of temperature (positive and negative), main effect of task (positive and negative) and interaction contrast (positive and negative).

Given the qualitative similarities and lack of demonstrable statistical differences we went ahead with our planned intention to combine the three datasets and all subsequent results relate to the *pooled cohort* comprising 57 subjects. We also note that the use of strict cluster thresholds for the brain, and of permutation testing for ROI-based analyses in ‘noisy’ brainstem regions, can produce robust and reproducible results even with a sample size of 20 (Brooks et al, 2017).

### Behavioural analysis (Pooled cohort)

The average high (noxious) temperature in the *pooled cohort* was 43.4°C (range 42°C - 45°C). Analysis of the pain ratings showed the expected main effect of temperature (F(1, 56) = 252.799, P < 0.0001, repeated measures ANOVA) and a task x temperature interaction (F(1, 56) = 31.969, P < 0.0001, Fig. 1). The main effect of task was not statistically significant (F(1,56) = 2.935, P = 0.092). A post-hoc paired t-test showed performance of the hard task produced a decrease in pain scores in the high temperature condition (mean *hard|high* = 38.1, SD 17.0 vs *easy|high* = 42.1, SD 16.5, P < 0.0001, Bonferroni corrected), consistent with an attentional analgesic effect (Fig 1).

**Figure 1.**
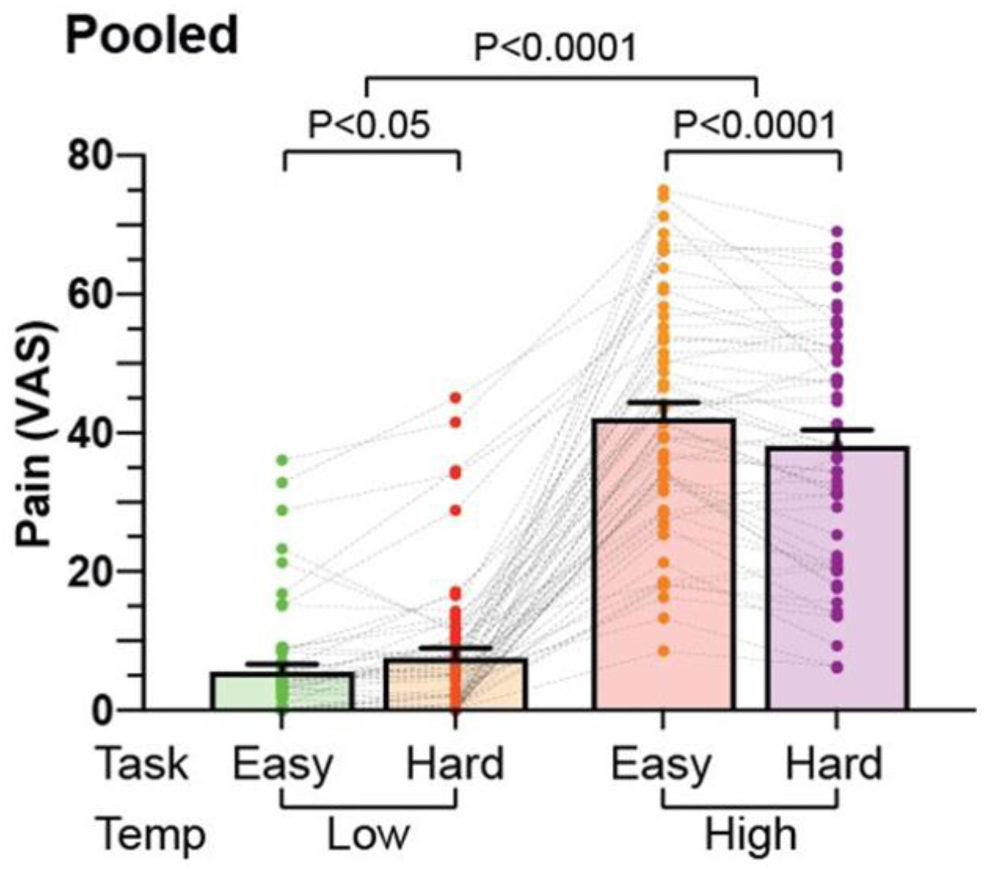
Pain ratings across experimental conditions (means with error bars representing the standard error of the mean) for the pooled cohort. A 2-way repeated measures ANOVA on the pain ratings showed the expected main effect of temperature (P < 0.0001) and a task x temperature interaction (P < 0.0001). A post-hoc paired t-test showed a decrease in pain scores in the high temperature condition during the hard task compared to the easy task (P < 0.0001), showing an analgesic effect caused by the increased cognitive load, as well as an increase in pain scores in the low temperature condition during the hard task compared to the easy task (P < 0.05). The main effect of task was not significant (P = 0.92). Error bars represent the standard error of the mean.

### Whole Brain & Brainstem-focussed analysis (Pooled cohort)

Activations were found for the positive main effect of temperature in a range of regions including the anterior and posterior cingulate cortices, precuneus, cerebellum, post-central gyrus (S1), dorsal posterior insula and opercular cortex, in the latter three cases with more prominent clusters contralateral to the side of thermal stimulation (Fig. 2A). In the negative main effect of temperature, significant clusters were found in the frontal medial cortex and in the subcallosal cortex (Fig 2A). At this whole brain level, no clusters of activity were found in the brainstem. To improve our ability to resolve activity in hindbrain structures, we undertook permutation testing using a *whole brainstem* mask which revealed clusters of activation in the positive main effect of temperature in the ventral PAG, LC bilaterally as well as the RVM (Fig 3A, p < 0.05, TFCE corrected).

**Figure 2.**
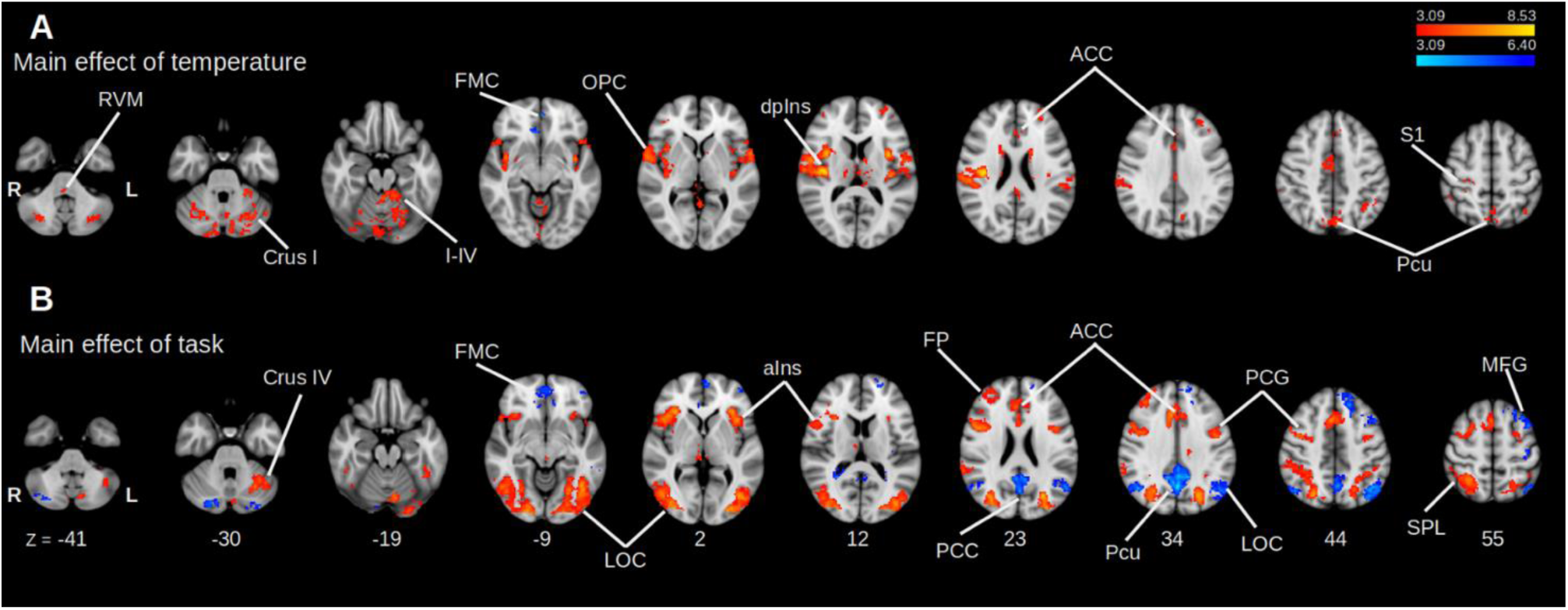
Main effect analyses in the *pooled cohort,* whole brain analysis. Positive (red/yellow) and negative (blue/light-blue). Data was obtained from cluster-based thresholding using an initial threshold of Z > 3.09 and FWE corrected p < 0.05, one-sample t-test. **(A)** Main effect of temperature. Positive activation in the high temperature conditions was found in anterior cingulate cortex (ACC), thalamus (THAL), dorsal posterior insula (dpIns), precuneus (PCu), primary somatosensory cortex (S1) and rostroventromedial medulla (RVM). Activation in the negative main effect of temperature (low temperature vs high temperature) was observed only in the frontal medial cortex (FMC). **(B)** Main effect of **task**. Activity in the positive main effect was found in the anterior insula (aIns), lateral occipital cortex (LOC), ACC, superior parietal lobule (SPL). Activity in the negative main effect was found in the frontal pole (FP), posterior cingulate cortex (PCC) and Pcu.

**Figure 3.**
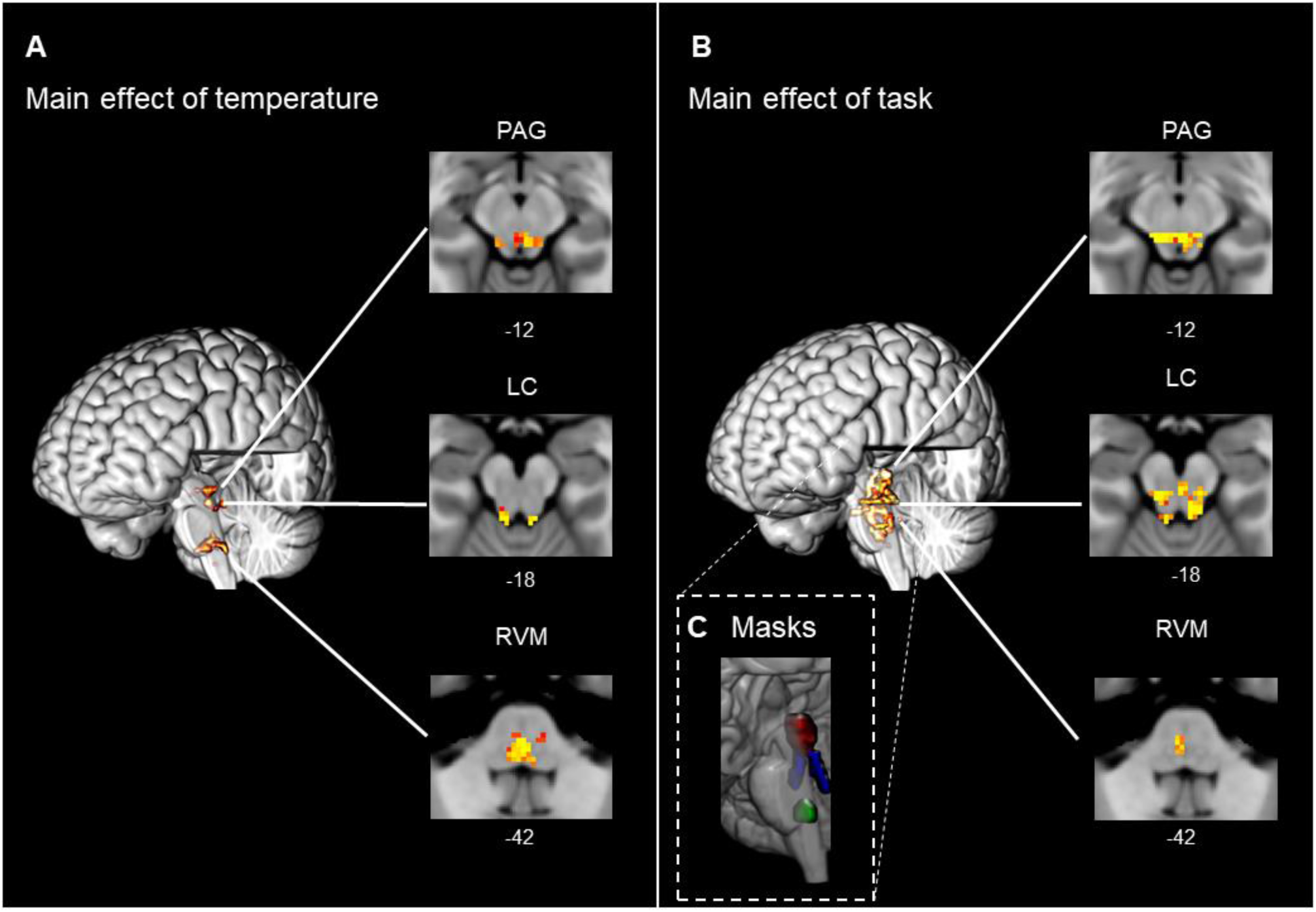
Main effect analyses in the brainstem. Results obtained after permutation testing with a probabilistic *whole brainstem* mask (p < 0.05, TFCE corrected). **(A)** Clusters of activation in the brainstem corresponding to the main effect of **temperature** was found in the ventral periaqueductal grey (PAG), rostroventromedial medulla (RVM) and bilateral locus coeruleus (LC). **(B)** Many areas in the superior brainstem show activity in the main effect of **task**, including the PAG, RVM and bilateral LC. **(C)** Anatomical masks defined in (Brooks et al., 2017), identifying the PAG (red), LC (blue), RVM (green).

Analysis of the positive main effect of task, showed extensive areas of activation within the lateral occipital cortex, superior parietal lobule, anterior cingulate cortex and anterior insula, as well as the PAG (Fig 2B). In the negative main effect of task, clusters were located in the posterior cingulate cortex, frontal medial cortex and in the lateral occipital cortex (Fig 2B). Permutation tests within the *whole brainstem* mask showed multiple clusters of activation, including in the LC bilaterally, RVM and PAG (Fig 3B, p < 0.05, TFCE corrected).

In the interaction contrast between task and temperature no cluster reached significance either at the whole brain level nor when using the whole brainstem masked analysis.

These findings from the *pooled cohort* showed close similarity to those of Brooks et al. (2017) with the same areas found in the main effects analysis (Table 1 shows all significant clusters). The additional findings at a whole brain level were that both the RVM and the precuneus now appear in the main effect of temperature and the dorsolateral PAG in the main effect of task (the RVM and PAG were only seen in a nucleus specific masked analysis in Brooks et al 2017). Similarly, activity in the brainstem is now seen in more areas using a whole brainstem mask rather than only in the nucleus specific masks (e.g. main effect of temperature in RVM alone previously versus RVM, LC and PAG in this *pooled* analysis).

**Table 1.**
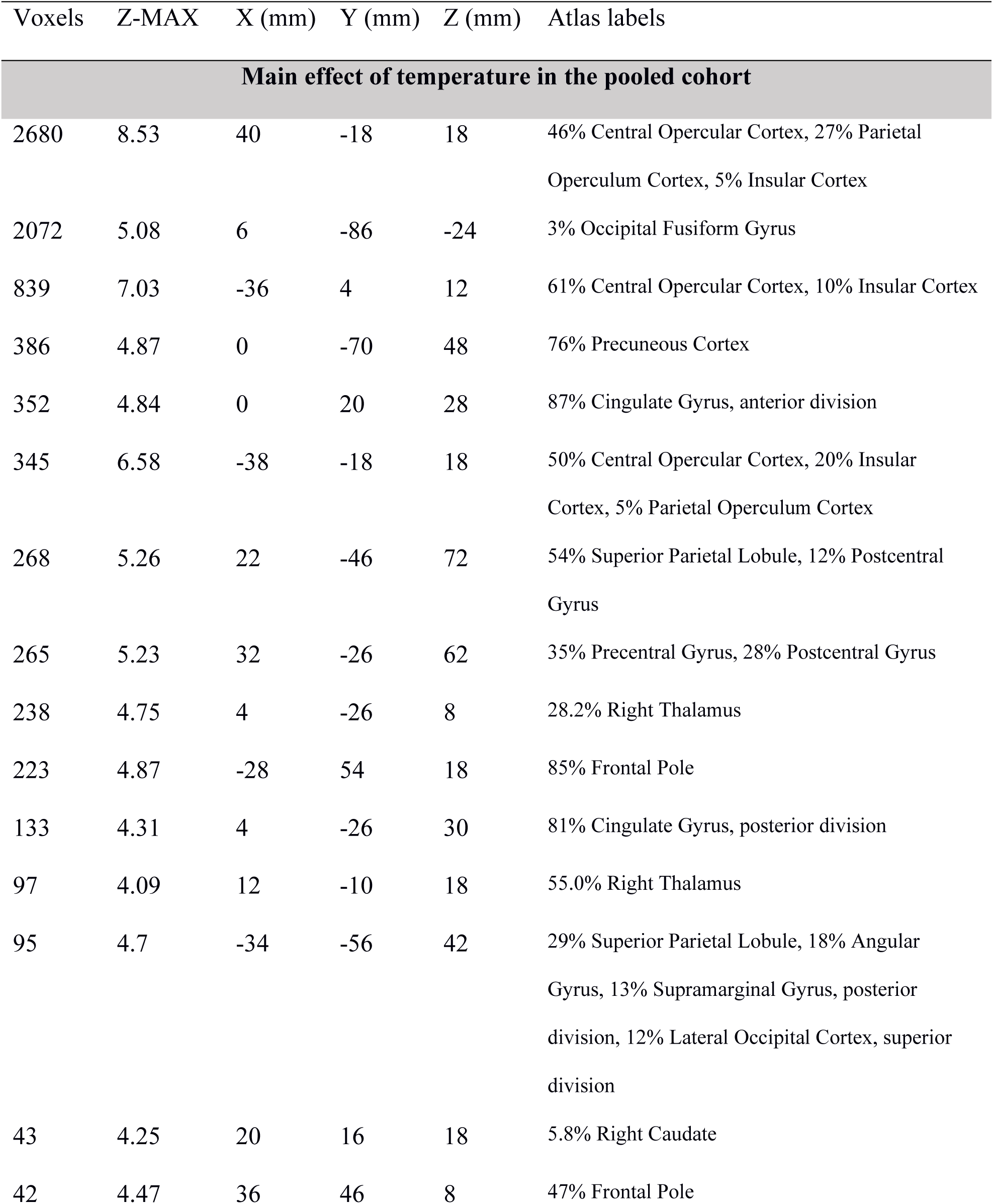
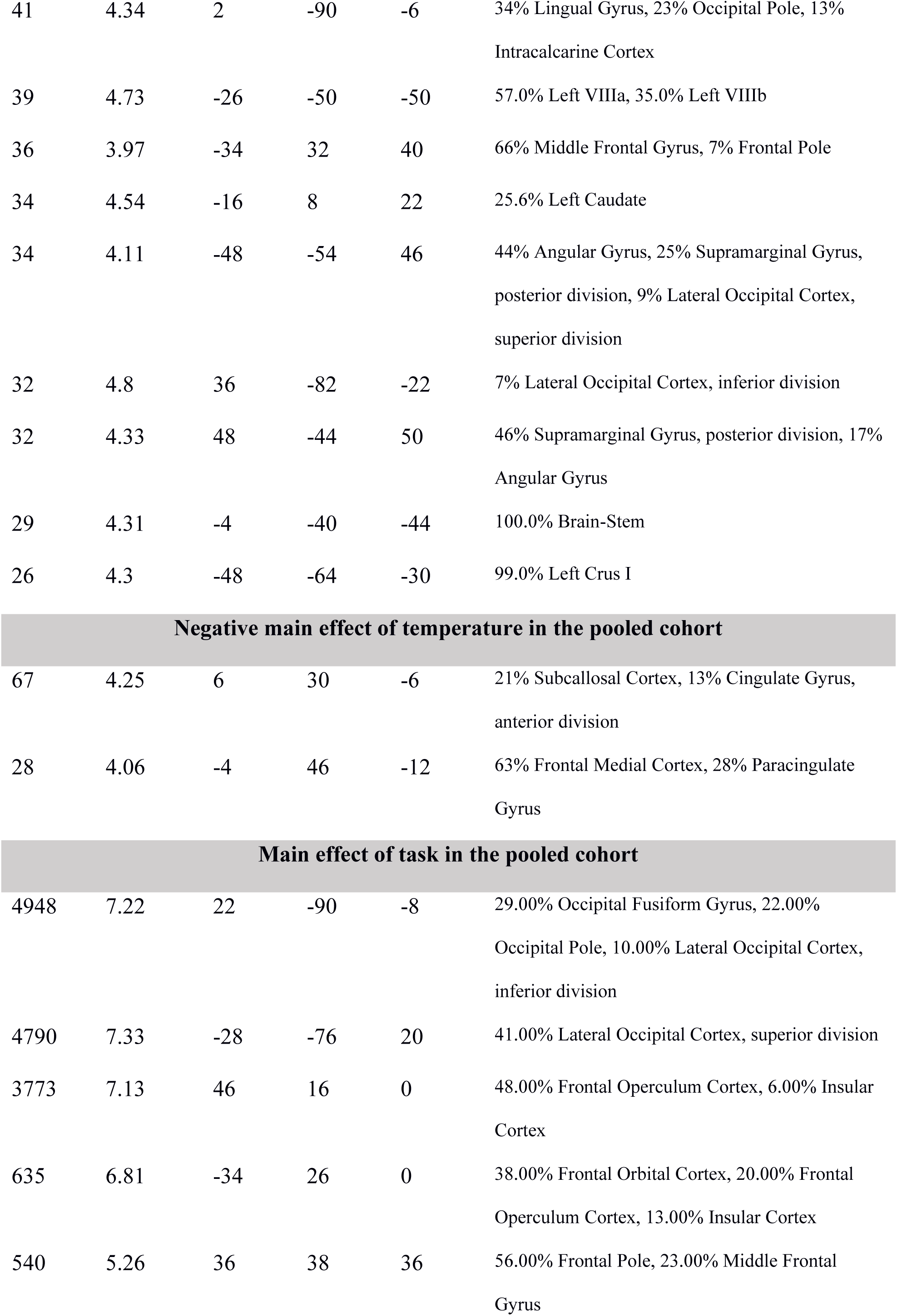
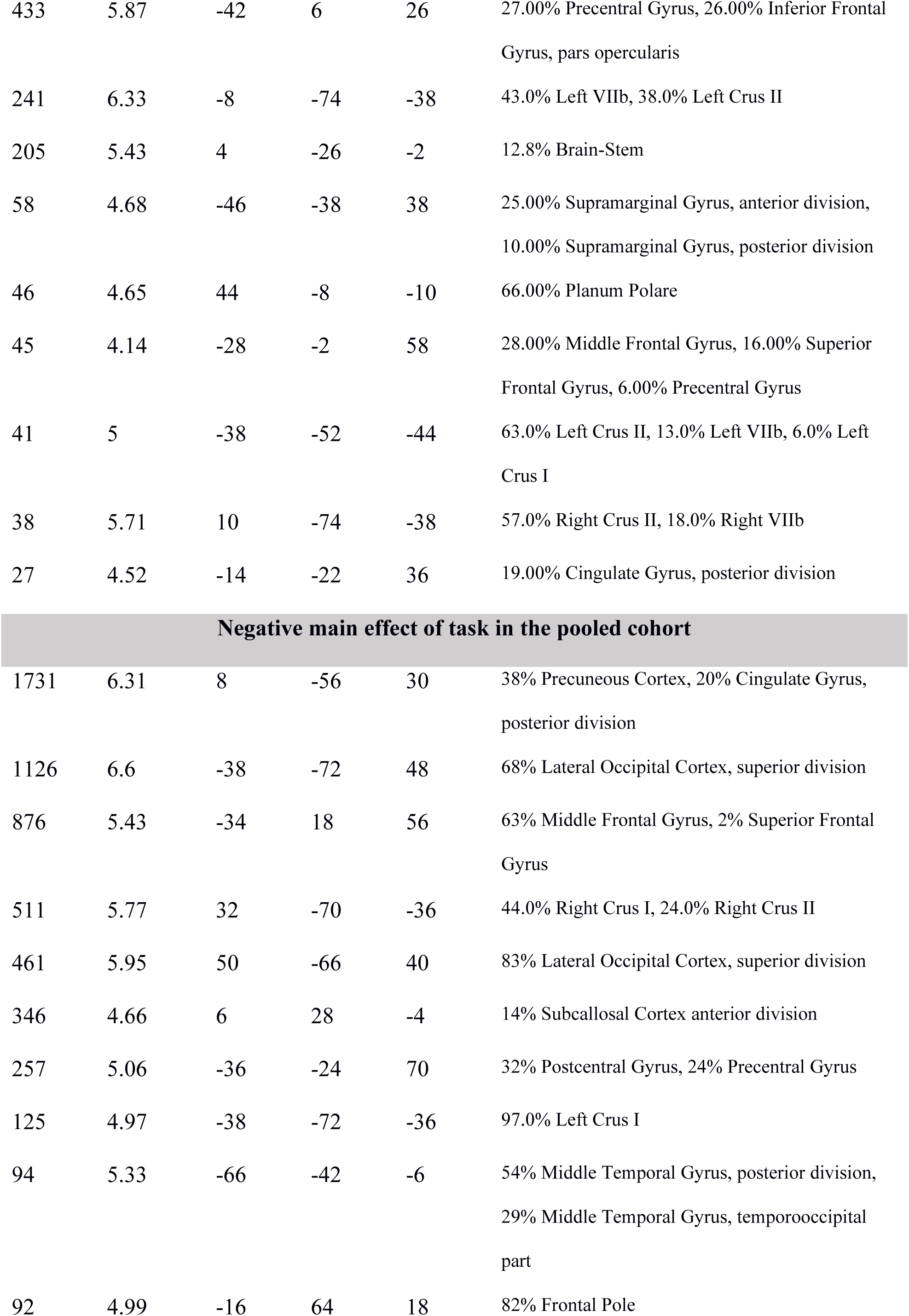
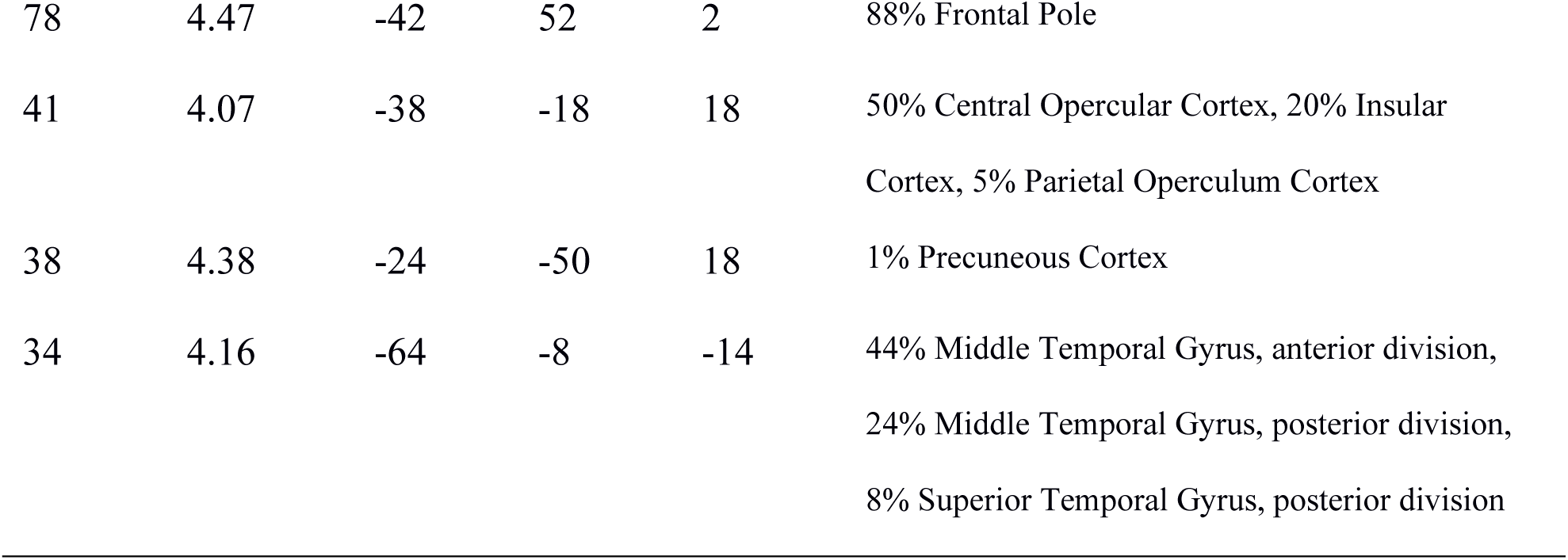
Activation clusters from main effects of temperature and distraction in the pooled cohort obtained with cluster-forming threshold Z>3.09 and cluster-corrected p<0.05. The tables were created with Autoaq (part of FSL), with atlas labels based on the degree of overlap with probabilistic atlases (Harvard Oxford Cortical Structural Atlas, Harvard Oxford Subcortical Structural Atlas, Cerebellar Atlas in MNI152 space after normalization with FNIRT). Only those structures to which the cluster had a >5% chance of belonging to are presented.

| Voxels | Z-MAX | X (mm) | Y (mm) | Z (mm) | Atlas labels |
| --- | --- | --- | --- | --- | --- |
| **Main effect of temperature in the pooled cohort** |  |  |  |  |  |
| 2680 | 8.53 | 40 | -18 | 18 | 46% Central Opercular Cortex, 27% Parietal Operculum Cortex, 5% Insular Cortex |
| 2072 | 5.08 | 6 | -86 | -24 | 3% Occipital Fusiform Gyrus |
| 839 | 7.03 | -36 | 4 | 12 | 61% Central Opercular Cortex, 10% Insular Cortex |
| 386 | 4.87 | 0 | -70 | 48 | 76% Precuneous Cortex |
| 352 | 4.84 | 0 | 20 | 28 | 87% Cingulate Gyrus, anterior division |
| 345 | 6.58 | -38 | -18 | 18 | 50% Central Opercular Cortex, 20% Insular Cortex, 5% Parietal Operculum Cortex |
| 268 | 5.26 | 22 | -46 | 72 | 54% Superior Parietal Lobule, 12% Postcentral Gyrus |
| 265 | 5.23 | 32 | -26 | 62 | 35% Precentral Gyrus, 28% Postcentral Gyrus |
| 238 | 4.75 | 4 | -26 | 8 | 28.2% Right Thalamus |
| 223 | 4.87 | -28 | 54 | 18 | 85% Frontal Pole |
| 133 | 4.31 | 4 | -26 | 30 | 81% Cingulate Gyrus, posterior division |
| 97 | 4.09 | 12 | -10 | 18 | 55.0% Right Thalamus |
| 95 | 4.7 | -34 | -56 | 42 | 29% Superior Parietal Lobule, 18% Angular Gyrus, 13% Supramarginal Gyrus, posterior division, 12% Lateral Occipital Cortex, superior division |
| 43 | 4.25 | 20 | 16 | 18 | 5.8% Right Caudate |
| 42 | 4.47 | 36 | 46 | 8 | 47% Frontal Pole |
| 41 | 4.34 | 2 | -90 | -6 | 34% Lingual Gyrus, 23% Occipital Pole, 13% Intracalcarine Cortex |
| 39 | 4.73 | -26 | -50 | -50 | 57.0% Left VIIla, 35.0% Left VIIlb |
| 36 | 3.97 | -34 | 32 | 40 | 66% Middle Frontal Gyrus, 7% Frontal Pole |
| 34 | 4.54 | -16 | 8 | 22 | 25.6% Left Caudate |
| 34 | 4.11 | -48 | -54 | 46 | 44% Angular Gyrus, 25% Supramarginal Gyrus, posterior division, 9% Lateral Occipital Cortex, superior division |
| 32 | 4.8 | 36 | -82 | -22 | 7% Lateral Occipital Cortex, inferior division |
| 32 | 4.33 | 48 | -44 | 50 | 46% Supramarginal Gyrus, posterior division, 17% Angular Gyrus |
| 29 | 4.31 | -4 | -40 | -44 | 100.0% Brain-Stem |
| 26 | 4.3 | -48 | -64 | -30 | 99.0% Left Crus I |
| Negative main effect of temperature in the pooled cohort |  |  |  |  |  |
| 67 | 4.25 | 6 | 30 | -6 | 21% Subcallosal Cortex, 13% Cingulate Gyrus, anterior division |
| 28 | 4.06 | -4 | 46 | -12 | 63% Frontal Medial Cortex, 28% Paracingulate Gyrus |
| Main effect of task in the pooled cohort |  |  |  |  |  |
| 4948 | 7.22 | 22 | -90 | -8 | 29.00% Occipital Fusiform Gyrus, 22.00% Occipital Pole, 10.00% Lateral Occipital Cortex, inferior division |
| 4790 | 7.33 | -28 | -76 | 20 | 41.00% Lateral Occipital Cortex, superior division |
| 3773 | 7.13 | 46 | 16 | 0 | 48.00% Frontal Operculum Cortex, 6.00% Insular Cortex |
| 635 | 6.81 | -34 | 26 | 0 | 38.00% Frontal Orbital Cortex, 20.00% Frontal Operculum Cortex, 13.00% Insular Cortex |
| 540 | 5.26 | 36 | 38 | 36 | 56.00% Frontal Pole, 23.00% Middle Frontal Gyrus |
| 433 | 5.87 | -42 | 6 | 26 | 27.00% Precentral Gyrus, 26.00% Inferior Frontal Gyrus, pars opercularis |
| 241 | 6.33 | -8 | -74 | -38 | 43.0% Left VIIb, 38.0% Left Crus II |
| 205 | 5.43 | 4 | -26 | -2 | 12.8% Brain-Stem |
| 58 | 4.68 | -46 | -38 | 38 | 25.00% Supramarginal Gyrus, anterior division, 10.00% Supramarginal Gyrus, posterior division |
| 46 | 4.65 | 44 | -8 | -10 | 66.00% Planum Polare |
| 45 | 4.14 | -28 | -2 | 58 | 28.00% Middle Frontal Gyrus, 16.00% Superior Frontal Gyrus, 6.00% Precentral Gyrus |
| 41 | 5 | -38 | -52 | -44 | 63.0% Left Crus II, 13.0% Left VIIb, 6.0% Left Crus I |
| 38 | 5.71 | 10 | -74 | -38 | 57.0% Right Crus II, 18.0% Right VIIb |
| 27 | 4.52 | -14 | -22 | 36 | 19.00% Cingulate Gyrus, posterior division |
| Negative main effect of task in the pooled cohort |  |  |  |  |  |
| 1731 | 6.31 | 8 | -56 | 30 | 38% Precuneous Cortex, 20% Cingulate Gyrus, posterior division |
| 1126 | 6.6 | -38 | -72 | 48 | 68% Lateral Occipital Cortex, superior division |
| 876 | 5.43 | -34 | 18 | 56 | 63% Middle Frontal Gyrus, 2% Superior Frontal Gyrus |
| 511 | 5.77 | 32 | -70 | -36 | 44.0% Right Crus I, 24.0% Right Crus II |
| 461 | 5.95 | 50 | -66 | 40 | 83% Lateral Occipital Cortex, superior division |
| 346 | 4.66 | 6 | 28 | -4 | 14% Subcallosal Cortex anterior division |
| 257 | 5.06 | -36 | -24 | 70 | 32% Postcentral Gyrus, 24% Precentral Gyrus |
| 125 | 4.97 | -38 | -72 | -36 | 97.0% Left Crus I |
| 94 | 5.33 | -66 | -42 | -6 | 54% Middle Temporal Gyrus, posterior division, 29% Middle Temporal Gyrus, temporooccipital part |
| 92 | 4.99 | -16 | 64 | 18 | 82% Frontal Pole |
| 78 | 4.47 | -42 | 52 | 2 | 88% Frontal Pole |
| 41 | 4.07 | -38 | -18 | 18 | 50% Central Opercular Cortex, 20% Insular Cortex, 5% Parietal Operculum Cortex |
| 38 | 4.38 | -24 | -50 | 18 | 1% Precuneous Cortex |
| 34 | 4.16 | -64 | -8 | -14 | 44% Middle Temporal Gyrus, anterior division, 24% Middle Temporal Gyrus, posterior division, 8% Superior Temporal Gyrus, posterior division |

### Linear pain encoding regions

Brain regions whose activity was linearly related to perceived pain intensity were identified using an *intra*-subject parametric regression. This revealed a network of positively correlated regions (similar to those seen in the main effect of temperature) including primarily the right (contralateral) dorsal posterior insula and S1, the anterior cingulate cortex, frontal lobe and the precuneus (Fig 4A). Regions showing a linear decrease in activation with pain ratings were restricted to the occipital cortex bilaterally and ipsilateral primary somatosensory cortex (Fig 4A). Permutation testing in the brainstem (using RVM, PAG and LC masks) identified only the RVM as showing a positive correlation with pain intensity (Fig 4B). No region showed a negative correlation with pain. All these findings were consistent with Brooks *et al* (2017), with the addition of a cluster identified in the thalamus (Table 2).

**Figure 4.**
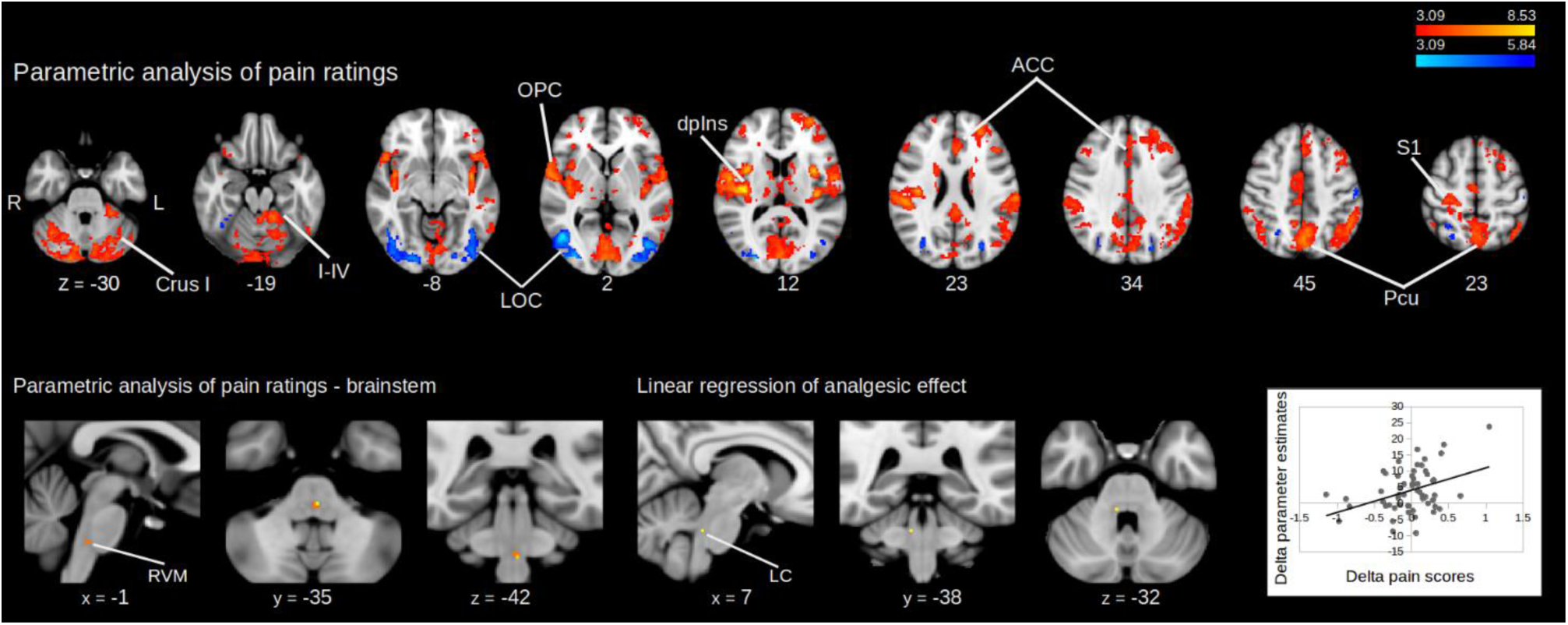
**A)** Intrasubject parametric regression with pain ratings across all the experimental conditions, in the whole brain. In red-yellow are shown regions whose activity linearly increase with pain ratings, in blue-light blue regions whose activity decreases with the pain ratings. (height threshold Z > 3.09, corrected cluster extent threshold p < 0.05) **B)** Intrasubject parametric regression with pain ratings, using RVM, LC and PAG masks. Only the RVM shows linear increase in activity with the pain scores (p < 0.05, TFCE corrected). **C)** Inter-subject parametric regression with the analgesic effect (i.e. ratings of *easy|high* – *hard|high*), using a LC mask (p < 0.05, TFCE corrected).

**Table 2.** Results from intrasubject parametric regression with pain ratings in the *pooled cohort* obtained with cluster-forming threshold Z>3.09 and cluster-corrected p<0.05. The tables were created with Autoaq (part of FSL), with atlas labels based on the degree of overlap with probabilistic atlases (Harvard Oxford Cortical Structural Atlas, Harvard Oxford Subcortical Structural Atlas, Cerebellar Atlas in MNI152 space after normalization with FNIRT). Only those structures to which the cluster had a >5% chance of belonging to are presented.

| Voxels | Z-MAX | X (mm) | Y (mm) | Z (mm) | Atlas labels |
| --- | --- | --- | --- | --- | --- |
| **Positive relationship with pain ratings** |  |  |  |  |  |
| 9467 | 6.36 | 32 | -28 | 64 | 40% Postcentral Gyrus, 26% Precentral Gyrus |
| 3621 | 8.37 | 40 | -18 | 18 | 46% Central Opercular Cortex, 27% Parietal Operculum Cortex, 5% Insular Cortex |
| 3481 | 7.78 | -56 | -2 | 8 | 45% Central Opercular Cortex, 28% Precentral Gyrus, 5% Planum Polare |
| 1693 | 5.93 | -26 | 46 | 26 | 79% Frontal Pole |
| 287 | 4.98 | -62 | -56 | -10 | 54% Middle Temporal Gyrus, temporooccipital part, 21% Inferior Temporal Gyrus, temporooccipital part, 7% Lateral Occipital Cortex, inferior division |
| 285 | 5.2 | 4 | -26 | 8 | 28.2% Right Thalamus |
| 120 | 4.42 | 20 | -8 | 28 | 3.1% Right Caudate |
| 95 | 4.18 | 42 | 48 | 8 | 80% Frontal Pole |
| 86 | 4.71 | 16 | -20 | 10 | 099.8% Right Thalamus |
| 71 | 5.79 | -26 | -50 | -50 | 57% Left VIIla, 35% Left VIIlb |
| 59 | 4.41 | 46 | 26 | 36 | 65% Middle Frontal Gyrus |
| 46 | 3.82 | -20 | -28 | 68 | 39% Precentral Gyrus, 23% Postcentral Gyrus |
| 38 | 4.17 | -42 | -72 | 24 | 72% Lateral Occipital Cortex, superior division |
| 37 | 4.11 | -18 | -38 | 66 | 51% Postcentral Gyrus |
| 36 | 3.84 | 2 | -66 | -38 | 69% Vermis VIIla, 13% Vermis VIIlb |
| 34 | 3.83 | 24 | 54 | 26 | 76% Frontal Pole |
| 33 | 4.48 | 52 | 30 | 22 | 32% Middle Frontal Gyrus, 30% Inferior Frontal Gyrus, pars triangularis |
| 31 | 3.78 | 28 | 58 | 0 | 76% Frontal Pole |
| Negative relationship with pain ratings |  |  |  |  |  |
| 1637 | 6.16 | 46 | -64 | 2 | 56% Lateral Occipital Cortex, inferior division,<br>10% Middle Temporal Gyrus, temporooccipital<br>part |
| 1184 | 5.36 | -42 | -74 | 0 | 63% Lateral Occipital Cortex, inferior division |
| 176 | 5.38 | -26 | -76 | 30 | 67% Lateral Occipital Cortex, superior division |
| 126 | 4.73 | 26 | -52 | 50 | 34% Superior Parietal Lobule, 3% Lateral Occipital<br>Cortex, superior division |
| 51 | 4.32 | -54 | -16 | 52 | 57% Postcentral Gyrus, 8% Precentral Gyrus |

### Regions whose activity correlates with analgesic effect

An *inter*-subject whole-brain mixed effects comparison between the *hard|high* and *easy|high* conditions did not identify any region whose activity linearly correlated with the differences in pain ratings (i.e. analgesia). A parametric regression showed a linear relationship between activity and analgesic effect only in the contralateral (right) LC (i.e. decreased pain ratings were associated with increased BOLD), after permutation testing with LC, RVM and PAG masks (Fig. 4C). A positive relationship was noted between the parameter estimates extracted from the rLC and the attentional analgesic effect on pain scores (Fig 4C).

### gPPI analysis

The analysis aimed to identify changes in effective connectivity associated with altered task difficulty, temperature and the task x temperature interaction. The previous main effect analyses provided us with functional activation masks that were used to extract time courses for these gPPI analyses; in the RVM for the main effect of temperature, and in the PAG, rLC and ACC for the main effect of task (see Methods). Permutation testing revealed increased connectivity with the following contrasts (Fig 5):

- RVM seed - increased connectivity to PAG for the interaction contras
- ACC seed - increased connectivity with the right (contralateral) LC in the interaction contrast and with the PAG in the main effect of task
- PAG seed - did not show any significant change in effective connectivity
- rLC seed - did not show any significant change in effective connectivity.

**Figure 5.**
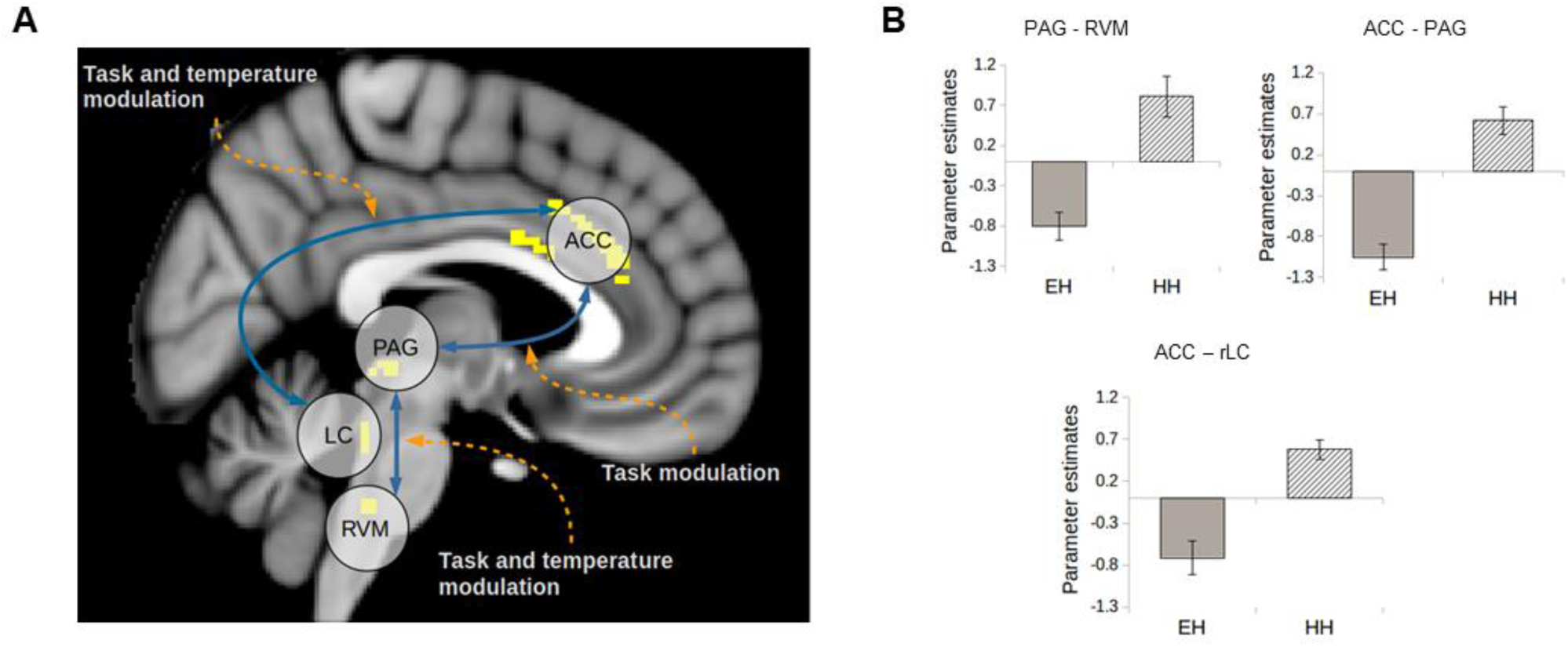
**(A)** Schematic representation of results of the PPI analysis. Results were obtained with single-nucleus functional masks and permutation testing (P < 0.05, TFCE corrected). **(B)** Parameter estimates extracted from the peak destination voxel from the PPI analysis (see text for details), in the *easy|high* and *hard|high* conditions.

For all gPPI results, parameter estimates were extracted from the voxel with greatest significance in each individual to explore the nature of these interactions (Fig 5B). In all cases, the parameter estimates were greater in the *hard|high* compared to the *easy|high* condition, indicating an increase in coupling in the circumstances producing analgesia.

### DCM

The connections resolved in the gPPI were further examined with DCM to resolve the directionality of the task effect. We systematically varied the location of the task inputs and modulation, while the temperature modulation was kept fixed in all models as a bottom-up effect. External inputs were both hard/easy task and high/low temperature, while modulations were only hard task and high temperature. Five models were specified (Fig 6A):

1. No modulation of connections,
2. Task was bottom-up on the ACC-PAG-RVM and on the ACC-LC axis.
3. Task was top-down for both pathways.
4. Task was top-down in the ACC-PAG-RVM axis and bottom-up in the ACC-LC connection.
5. Task was bottom-up in ACC-PAG-RVM and top-down in ACC-LC.

**Figure 6.**
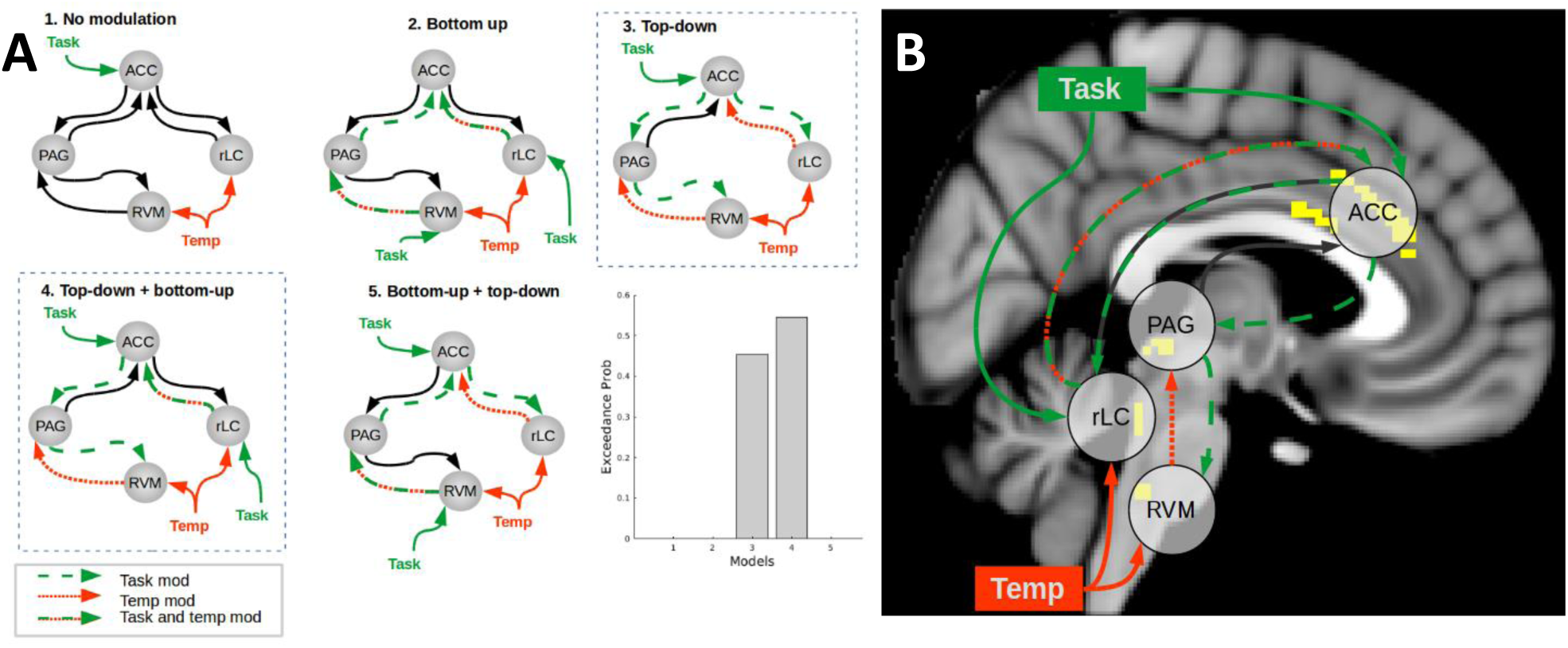
(A) Specified models and result of Bayesian Model Selection. The effect of temperature is always bottom-up, while the task could have a bottom-up or top-down effect. Model 3 and 4 have the strongest evidence of reproducing the data, with slightly stronger evidence for model 4. (B) Schematic representation of interactions during attentional analgesia, after PPI and DCM.

Models 3 and 4 were found to best fit the data in BMS, with protected exceedance probability = 0.45 and 0.55 respectively (Fig 6B), and Bayesian omnibus risk of zero. In both, the task had a top-down influence on ACC-PAG-RVM, while the ACC-LC connection was top-down modulated in one model and bottom-up modulated in the other. Bayesian model averaging was used to extract parameter estimates (Table 3).

**Table 3.** Summary of mean parameter estimates for connections and task modulation in DCM.

| Parameter | Parameter estimate mean (SD) | Task modulation mean (SD) |
| --- | --- | --- |
| ACC-PAG connection | 0.084 (0.0012) | 0.029 (0.0052) |
| ACC-LC connection | 0.058 (0.0011) | -0.019 (0.0054) |
| PAG-ACC connection | 0.083 (0.0012) | -0.0058 (0.0026) |
| PAG-RVM connection | 0.039 (0.0011) | -0.038 (0.0061) |
| LC-ACC connection | 0.054 (0.0012) | -0.012 (0.0052) |
| RVM-PAG connection | 0.034 (0.0012) | -0.0004 (0.0026) |

All connections were also tested with an analgesic covariate, to find whether one or more consistently differed in participants that showed an analgesic effect. No connection reached significance in this test.

## Discussion

In the context of a reproducibility crisis that is afflicting neuroscience, especially in fMRI experiments (Button et al., 2013; Eklund et al., 2016), we report results that have reproduced and extended the generalisability of the key findings regarding the neural substrates of attentional analgesia that were originally reported in Brooks et al., 2017. The same subset of brain regions was shown to be involved in the performance of a visual attention task and in pain perception. Importantly, we replicated the involvement in attentional analgesia of brainstem structures such as PAG, LC and RVM. This was demonstrable without the need for region specific anatomical masks because we increased our resolving power by analysing a larger cohort of subjects. By using gPPI and DCM analyses we demonstrated changes in effective connectivity between these key brainstem regions and the ACC, that are likely involved in the mediation of attentional analgesia.

The higher statistical power provided by 57 subjects, some 3-fold greater than in Brooks et al. (2017), yielded stronger findings especially in the brainstem nuclei. For the main effect of temperature, we were now able to see activation in the RVM at a whole brain analysis level. Additionally, using a full brainstem mask (rather than region specific masks) we found activation not only in the RVM, as previously (Brooks et al., 2017), but also in PAG and bilaterally in LC. While it has long been known from animal studies that these brainstem regions receive nociceptive input from the spinal cord (Blomqvist & Craig, 1991; Cedarbaum & Aghajanian, 1978; Keay et al., 1997) this has seldom been clearly demonstrated in human imaging studies. The specificity of this pattern of nociceptive information flow is striking with activations confined to discrete territories including ventral PAG, LC and RVM as well as activations in the region of parabrachial nucleus, nucleus solitarius, sub nucleus reticularis dorsalis and nucleus cuneiformis. The intra-subject linear regression analysis with pain scores in regions such as the dorsal posterior insula and the anterior cingulate cortex, where the BOLD signal linearly scales with perceived pain, was also consistent with previous results (Horing et al., 2019). In the brainstem, the only region showing the same linear relationship is the RVM. This clearly demonstrates that this brainstem territory is likely to be playing an important role in coding nociceptive intensity. To our knowledge, no other single study has been able to produce such a complete activation map in the human brainstem in response to noxious stimulation (see review by Henderson & Keay, 2018).

Considering the main effect of task, we were now able to identify activity in the PAG at the whole brain level. By using a whole brainstem mask, we were able to detect activity in RVM and LC bilaterally (as well as the dorsal PAG), while in our previous study using region specific masks we could only find responses in the right LC and PAG (Brooks et al., 2017). It is interesting to note that the attention task recruited the dorsal and ventral PAG whereas the noxious input produced activation in the ventral region of the nucleus perhaps in line with the known behavioural specialisation of columns within this crucial integrating nucleus (Linnman et al., 2012; Roy et al., 2014).

The magnitude of the analgesic effect showed a correlation with activity in the right LC (a finding that we previously noted in Brooks et al. (2017) but was just below formal statistical significance). This was the only location in the neuroaxis that showed this relationship. The LC is well positioned both anatomically and functionally to mediate a component of attentional analgesia, not only because it is responsive to attentional states and cognitive task performance (Aston-Jones & Cohen, 2005; Sales et al., 2019; Sara, 2009; Yu & Dayan, 2005) and to nociceptive inputs (Cedarbaum & Aghajanian, 1978; Howorth et al., 2009), but also particularly in being able to cause analgesia via its direct spinal cord projections (Hirschberg et al., 2017; Jones et al., 1986). Intriguingly, the spinal cord-projecting neurons are located in the caudal part of the LC in rodents (Hirschberg et al., 2017), as is the LC region that we found to correlate with the analgesic effect. Previous analyses have demonstrated linear relationships between the analgesic effect and activity located in the PAG in 9 subjects (Tracey et al., 2002) and RVM in 20 subjects (Brooks et al., 2017). We note that neither of these findings were replicated our current study of 57 subjects. While all three findings are biologically plausible, a large sample size seems necessary to produce robust results with inter-subject regression (especially in small, noisy brainstem nuclei) and this is likely to be complicated by the known interactions between these regions in nociceptive processing (discussed below).

To examine interplay between the cortical and brainstem structures we hypothesised to be involved in attentional analgesia, we performed a generalised PPI, which determines how connectivity changes as a result of experimental manipulation (i.e. effective connectivity). Our prior hypothesis for a role of the ACC was supported by the observation that it showed overlapping areas of activation in both conditions. We observed altered connectivity between the ACC and contralateral (right) LC during the interaction between task and temperature. Parameter estimates extracted from voxels showing this interaction revealed that coupling was enhanced during the hard task/high temperature condition. Furthermore, coupling increased between ACC and PAG with task demand, and between PAG and RVM, during the task x temperature interaction. Extraction of parameter estimates revealed that, as for ACC-LC connection, this PAG-RVM interaction was enhanced in the hard task/high temperature condition.

Effective connectivity changes in these pathways may therefore mediate the process of attentional analgesia. This could be achieved through LC projections to the ACC increasing the signal-to-noise (or salience) of one input over another (Manella et al., 2017; Muller et al., 2019; Sales et al., 2019; Sara, 1985; Vazey et al., 2018) and/or ACC to spinally projecting LC neurons modulating the activity of dorsal horn neurons (i.e. decreasing nociceptive transmission) both actions potentially giving ‘precedence’ to the task. One intriguing aspect of this interaction is the lateralised nature of relationship between the right LC and the analgesic effect (i.e. contralateral to the stimulus) - a finding that has previously been noted in rodent studies where noxious stimuli increase the activity in the contralateral LC to a greater effect (Cedarbaum et al., 1978). Similarly, the reduction in perceived pain could equally be via ACC recruiting the PAG and RVM to produce antinociception at a spinal level during the demanding attentional task (Millan, 2002). This conceptually extends previous studies that have identified the ACC-PAG connection as being involved in a distraction from pain (attentional analgesia) paradigm (Valet et al., 2004), as well as in a placebo analgesia paradigm (Petrovic, 2002). The PAG-RVM descending control system has also already been implicated in placebo analgesia (Eippert et al., 2009; Grahl et al., 2018) via an opioid-dependent mechanism. The behavioural component of attentional analgesia has been reported to be impaired by opioid blockade, possibly by disrupting connections between the ACC-PAG-RVM descending control system (Sprenger et al., 2012).

We used dynamic causal modelling to resolve the directionality of the connection changes that were shown in the gPPI, with the objective of better characterising the network mechanism generating analgesia. We employed stochastic DCM, which allows for modelling of random neuronal noise in the system, to improve network resolution in brainstem areas significantly affected by physiological noise (Brooks et al., 2013). This routine was shown to improve the characterization of network structure and parameter inference over deterministic DCM (Daunizeau et al., 2012; Osório et al., 2015) and has been widely used in resting state and task-based fMRI studies since its release (Kahan et al., 2014; Ma et al., 2015, 2014; Ray et al., 2016; Zhang et al., 2015). Bayesian Model selection validated the results of the gPPI by excluding, for lack of evidence, a model where no connection was modulated by task. We resolved a top-down influence of task on the ACC-PAG and PAG-RVM connections, consistent with a descending pain modulatory system being involved in attentional analgesia (Sprenger et al., 2012). The ACC-LC pathway was however not resolved as clearly, with similar evidence in BMS for task modulation of the top-down and bottom-up connection. As discussed above, both possibilities are supported by solid biological evidence and it is possible that the two regions work in a feedback loop during attentional analgesia. On examination of the parameter estimates, it was noted that the task modulation had a negative effect on all connections, except for ACC-PAG. This is consistent with a reciprocal negative feedback loop between ACC and LC (Breton-Provencher et al., 2019; Ramos et al., 2007), and with a disinhibition of the RVM “off-cells” by the PAG (Lau et al., 2014). It is quite conceivable that the parallel ACC-LC and ACC-PAG-RVM systems described here work in concert to cause analgesia. Previous animal studies show that electrical stimulation of the PAG triggers noradrenaline release in the cerebrospinal fluid (Cui et al., 1999; Hammond et al., 1985) and the analgesic effect of stimulation can be partially blocked with intrathecal alpha2 antagonists.

We propose that the ACC acts to resolve the conflict caused by an attention-grabbing painful stimulus and the cognitive demands of a sustained visual attention task, by sending downstream signals to brainstem structures to facilitate optimal behaviour. Intriguingly, inputting the coordinates of the peak attentional activation of the ACC to Neurosynth (Yarkoni et al., 2011) identified three studies where the same region was involved in response to conflict (Barch et al., 2001; Scholl et al., 2017; van Veen et al., 2005; Wittfoth et al., 2008). In addition, voluntary control over the activation of this area was shown to result in modulation of pain perception in a neurofeedback study (deCharms et al., 2005). This network is also relevant for mindfulness-based techniques, where focus on an internal signal (e.g. breathing), effectively distracts subjects from the painful stimulus. Interestingly, mindfulness mediated pain relief is not mediated by endogenous opioids (Zeidan et al., 2016) but relies on the rACC (Zeidan et al., 2012; Zeidan & Vago, 2016), perhaps through the ACC-LC pathway.

In this study we have been able to resolve parallel cortical – brainstem pathways that form a network that is functionally engaged when pain perception is attenuated during attentional analgesia. We note that the spinal cord BOLD response to nociception has previously been shown to be modulated by attention (Sprenger et al., 2012). Whether this spinal modulation of nociception is the product of activation of the ACC-PAG-RVM and/or the ACC-LC system still needs to be demonstrated in humans. It is known that both pathways could involve opioids (Fields, 2004) and so previous studies using naloxone do not discriminate between these possibilities. A connectivity analysis examining the network activity between cortical territories, brainstem nuclei and dorsal horn *in toto* may help to define the key pathway in attentional analgesia. We further postulate that this network may be of importance in chronic conditions where disruption of attention and cognition (i.e. fibromyalgia) are co-morbid alongside pain.

